# Design of Multichannel Transcranial Temporal Interfering Stimulation System Using an Individual MRI

**DOI:** 10.64898/2025.12.25.696527

**Authors:** Sangkyu Bahn, Chany Lee, Bo-Yeong Kang

## Abstract

Transcranial temporal interfering stimulation (tTIS) is an electrical stimulation method, in which two high-frequency alternating electric fields generate interference in the deep brain. This study aimed to design and verify the performance of a system that can precisely stimulate the deep brain using a multichannel tTIS based on an MRI image of an individual’s brain. The optimization process, based on the parallel genetic algorithm with a GPU, was computationally verified using a modeled head that housed the deep brain. The hardware digitally stimulated the pre-interpreted head in a calculative manner, and the performance of the method was verified using an ideal head circuit. When four or more electrodes per frequency were used to stimulate the left thalamus, the rate of misstimulation was controlled to less than 2% on average for approximately 1 min. In the absence of deep-brain modeling, the average stimulus applied was reduced to 77.9%. The predicted signal had a 98.21% coefficient of determination for the modulations obtained by stimulating a head-type circuit using the proposed hardware. We designed a system that can stimulate the deep region of the brain through a multichannel tTIS, with due consideration for its practical application during the entire process.

CCS CONCEPTS: **• Theory and algorithms for application domains → Theory and algorithms for application domains; • Computing methodologies → Modeling and simulation; • Applied computing → Physical sciences and engineering**

## 1 INTRODUCTION

Neuromodulation is an alternative method for enhancing or treating brain function by stimulating the nervous system using electrical, magnetic, or chemical methods [32]. Transcranial electric stimulation (tES), a method of stimulating the brain by injecting a weak current through electrodes to generate electric fields, has been recognized as a safe and noninvasive method [37, 47, 50]. Grossmann proposed transcranial temporal interfering stimulation (tTIS) as a solution to stimulate the deep brain. This method causes interference in the deep brain using two high frequencies to which neurons do not respond [18]. Furthermore, numerous studies have been conducted on the physiological characteristics, optimization methods, and performance feasibility of this method [9, 19, 24, 34, 43, 55].

tTIS determines the distribution of interference modulation in the brain through a nonlinear combination of electrode currents with two types of frequencies. Therefore, determining the electrode currents that intensively stimulate the desired area is an extremely large and complex problem, and several studies have been conducted to optimize this problem [14, 23, 28, 42, 43, 54]. Recently, various methods have been proposed to increase the stimulation concentration using multiple channels [2, 23, 29]. One such method can automatically determine a local optimal solution in the solution space of the number of electrodes using an optimization algorithm for the placement of high-definition electrodes at the 10–10 system level [2, 23]. Although these studies demonstrated satisfactory outcomes at fixed electrode locations, they did not optimize electrode placement and naturally assumed deployment of a substantial number of electrodes across the scalp. However, practically applying the number of electrodes at the level of 10–10 systems to clinical practice is difficult [26], and a method for determining the most efficient arrangement of a small number of electrodes is often required. In a previous study, methods of manually adding a pair of electrodes to gradually produce better results [29] and using a genetic algorithm (GA) to optimize the positions of the electrodes were proposed [54, 58]. However, in multichannel optimization using combinations of multiple electrode signals, these methods require a significant computation time.

The methods for optimizing the interference stimulus were verified based on a realistic head model. A multilayered model with components such as the scalp, skull, cerebrospinal fluid (CSF), gray matter, and white matter has primarily been used to create a realistic head model from an MRI image [27]. This model is supported by various existing open-source tools [13, 22]. However, unlike traditional tES, which stimulates the shallow part of the brain, modeling of the complex structure inside the white matter is required to stimulate the deep brain. Previously, it was modeled manually [16]; however, with the development of deep learning, a network that can automatically segment detailed brain tissues has been developed [4, 20, 52]. However, multilayer models continue to be used in tTIS research based on the finite-element method (FEM) [2, 58].

In addition to computationally obtaining the value of the electrode current for the tTIS, implementing it in hardware is also important. Various commercial devices implementing tES have been developed [5, 35, 36]. For research purposes, isolators that output current proportional to the voltage of the function generator are widely used [3, 8, 10]. However, tTIS stimulation requires a current at the kilohertz level, and the existing commercial tES equipment does not support such high frequencies. Moreover, previous studies have reported that a commercial isolator, when connected to a function generator, does not operate normally because of distortion and attenuation problems at the kilohertz frequency level [10, 25, 31, 38]. To address these issues, a circuit capable of stimulating a current at the kilohertz level has been developed through a design that considers these issues in isolators, amplifiers, and electrodes [10, 11, 15].

Although tTIS research is actively progressing in various stages, it is focused on the characteristics and limitations of tTIS in specific fields, assuming that other processes are in the ideal state. In this study, a design for the entire system that considers the organic interaction at each stage, and their realistic application process in clinical situations is proposed. The specific methods proposed for the implementation of the system include the modeling method, which facilitates the utilization of deep segmentation results derived from deep learning in FEM-based optimization; the method to accelerate the computation of GA-based optimization; and the method to facilitate the straightforward and precise input of currents at kilohertz levels using the results of head model analysis. To validate the effectiveness of the system, we evaluated the impact of the presence or absence of deep modeling on the optimization results, efficiency of a small number of optimally located electrodes, computational time of the proposed method, and accuracy of the hardware created by the proposed method.

## 2 MATERIALS AND METHODS

### 2.1 Pipeline and scope of the tTIS system

Conventional tES methods stimulate near electrodes using a low frequency, which presents several difficulties when applied to tTIS. Recently, several issues have been addressed [2, 10, 11, 15, 23, 29]. However, these studies only verified the performance of a specific process under the assumption that other factors were ideal. For the practical application of the entire system, each component must be designed integrally.

For example, optimization studies of the electrode montage used a realistic head model consisting of a multilayered model with a uniform deep brain [23, 28]. This model is easy to implement and improves the optimization performance; however, it leads to differences between the predicted and actual results. In addition, several optimization studies, that consider only the calculation of the optimal current for a fixed electrode position, tend to use many electrodes, such as the 10–10 system. However, the number of electrodes used for optimization is directly related to hardware implementation, clinical preparation time, patient fatigue, and inaccuracies due to adherence [26]. The calculation time is also directly related to the waiting time experienced in a clinic. Similarly, hardware often requires expensive and heavy-function generators per channel for research purposes [11, 15], and increasing the hardware to implement multiple channels can result in several problems in clinical applications. In addition, the kilohertz acceptance of the stimulation device using the current source mentioned in Section 1 should be considered. As well as kilohertz acceptance, the isolator requires a very precise and noise-free voltage function generator to convert the voltage signal into a current signal as feedback in the circuit. Therefore, the entire system requires deep-brain modeling and optimization, including fast and efficient electrode placement as well as portable and programmable hardware that solves the problem of adapting tTIS to existing devices.

To overcome these issues, the proposed system deals with three subprocesses. The first is the process of obtaining segmentation results for the deep brain through deep learning and modeling it as a finite element with a polyhedral grid. The second is an optimization algorithm based on the parallel GA (PGA). This method accelerates the calculation speed through GPU operations by reconstructing the traditional GA by inducing a matrix operation. The last is a light and simple digital stimulation device that calculates and outputs a digital voltage signal, allowing the desired current to flow through a pre-interpreted head model.

In this paper, the detailed implementation of each method for applicable systems is described. In addition, we investigated the effects of the deep-brain model and efficiency of the GPU-based PGA on the optimization results using computer simulations based on FEM, as well as the hardware precision of the digital stimulator. Figure 1 shows the entire pipeline of the system for stimulating the desired area of the deep brain.

**Fig. 1.**
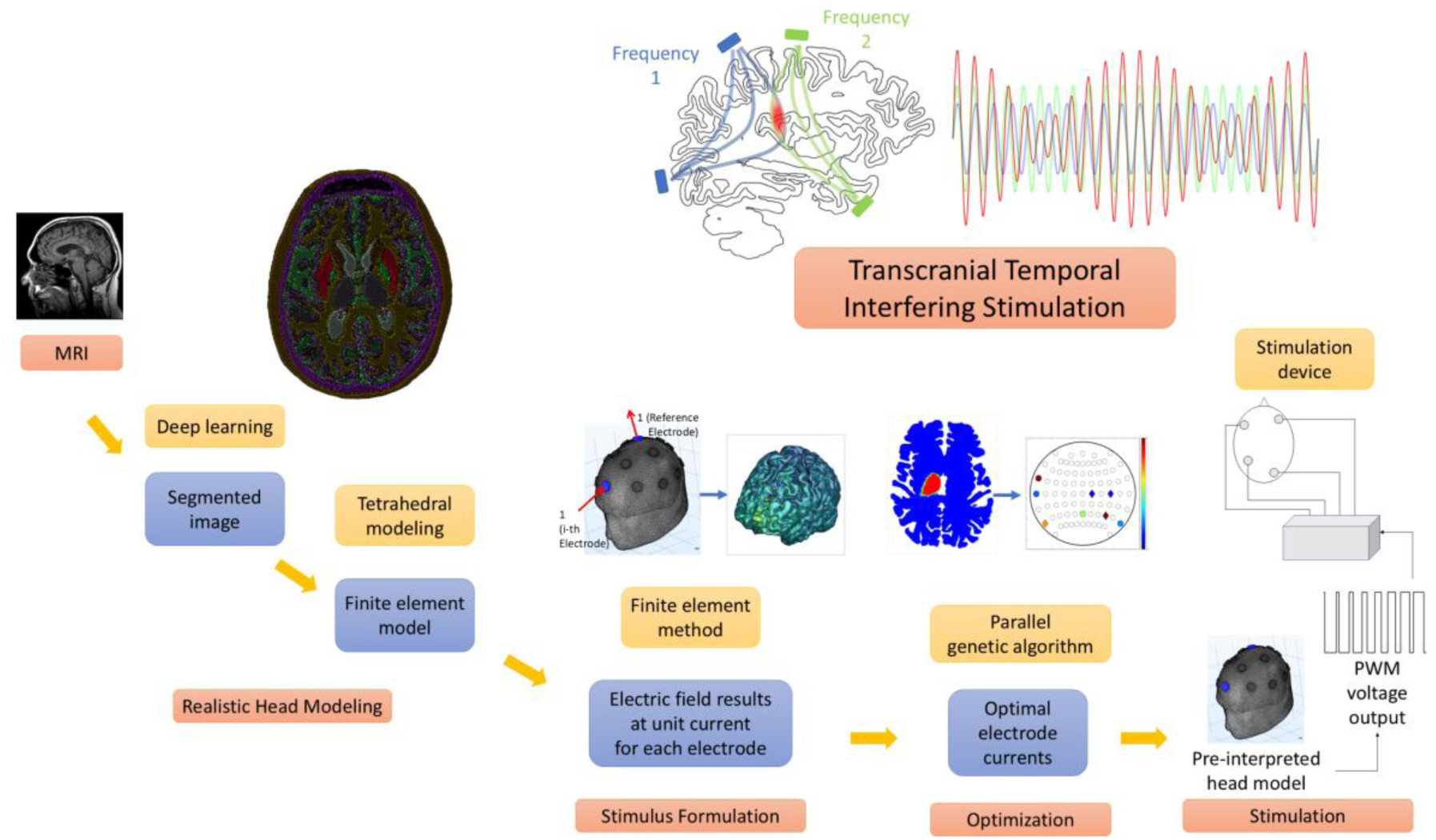
Overview of the tTIS and pipeline of the entire system for deep-brain stimulation from MRI images.

### 2.2 Realistic head model

Each tissue of the modeled head can be segmented using deep learning (particularly U-net [45]). Several public networks have been trained to rapidly obtain segmented images from MRI [4, 6, 20]. In this study, we propose a tetrahedral modeling method for FEM based on the premise of using deep learning for segmentation and analyze the effect of the presence or absence of deep-brain modeling. Therefore, we used the ground truth of the Benjamin network [4], which is the most granular segmentation data available, without considering the performance of the network itself.

Deep learning networks generate segmented 3D voxel images; however, for FEM, polyhedral finite-element modeling with smooth interfaces between tissues is required. A tetrahedron is the only polyhedron whose insides can be expressed by the linear interpolation of nodes in a 3D space, and its electromagnetic equation can be analytically derived without additional approximations, such as numerical integration. Furthermore, unlike hexahedrons, tetrahedrons have been primarily used in existing head modeling because of their advantage in flexibly adjusting the precision of the grid in areas that require shape precision. Therefore, this study proposes a tetrahedral modeling method that reduces the number of internal elements while maintaining the precision of the tissue surface using a voxel-image-based hexahedral model.

A voxel-segmented image was reconstructed into a tetrahedral finite-element model as follows. Each voxel can be modeled as a cube with six square faces (Figure 2(a)). The rectangular surface grid of the connected components in each organization was divided into two triangular surface grids (Figure 2(b)), and the angular grid was adjusted to obtain a smooth triangular grid surface using Laplacian smoothing [12] (Figure 2(c)). After designating the points inside the surface of the connected component for each tissue, a volumetric tetrahedral finite-element model was created using Delaunay triangulation [48] (Figure 2(d)). Laplacian smoothing and Delaunay triangulation were performed using the “Gibbon code” library [33]. The study was conducted on a single subject, and the modeled head consisted of 3,782,471 nodes with 23,023,778 tetrahedral elements. The sagittal cross-section of the finite-element model is shown in Figure 3(a), and the surface modeling results of the white matter are shown in Figure 3(b).

**Fig. 2.**
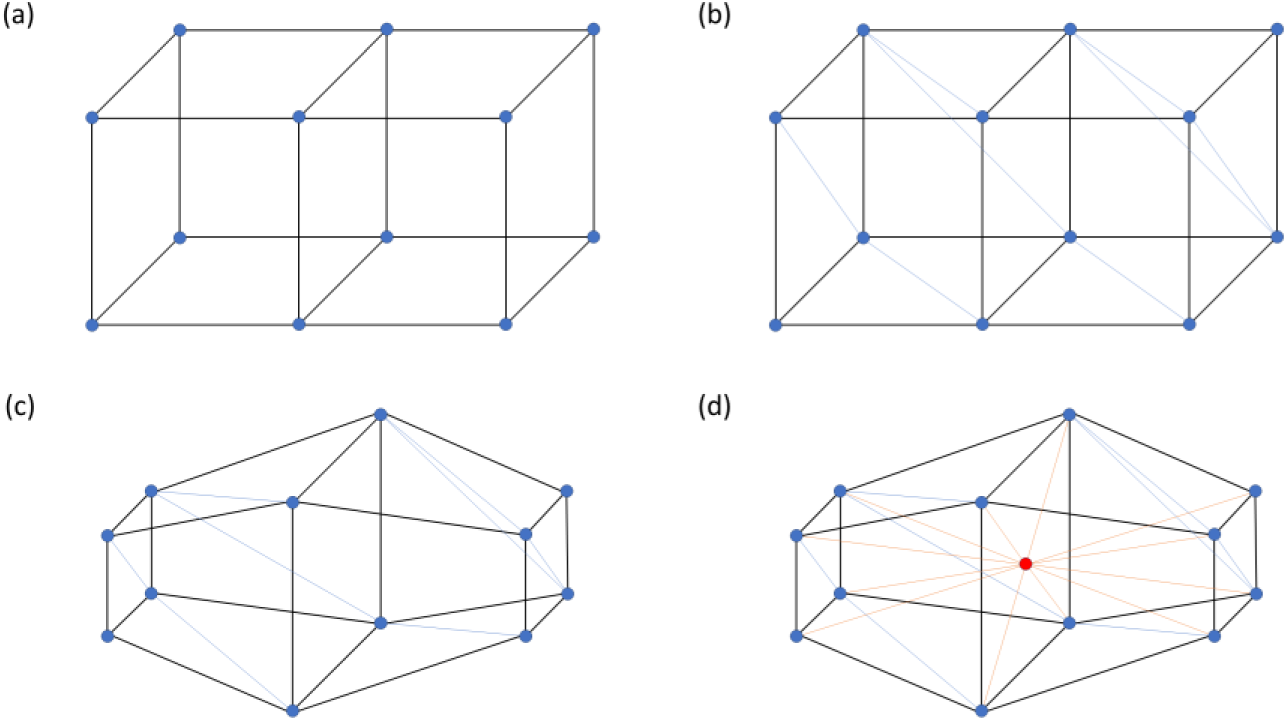
The process of tetrahedral finite-element modeling of a three-dimensional image. (a) A vowelized image is constituted by a combination of cubes. (b) The surface rectangle of each organization’s connected component is split into two triangles. (c) Laplacian smoothing is performed on the triangular surface grid of each connected component. (d) Create interpolation points inside the connected component and model the tetrahedron with a Delaunay triangulation.

**Fig. 3.**
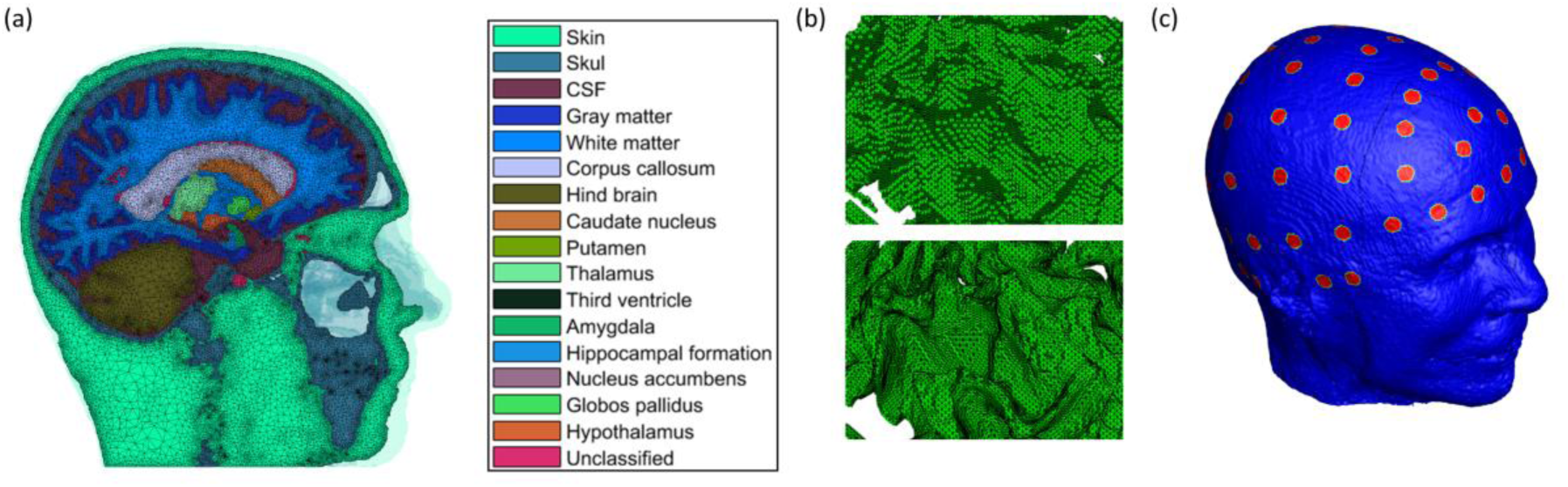
Realistic head modeling. (a) Sagittal cross-section of a finite-element model wherein the deep brain tissue is modeled. (b) Vowelized image of the gray matter surface and its tetrahedral finite-element modeling results. (c) Electrodes with a diameter of 1 cm arranged according to the international 10–10 system on a realistic head model.

The electrical conductivity of each tissue was specified based on previous tDCS studies [17], and the detailed tissue of the head was modeled and divided into 17 types (Figure 3(a), Table 1). In the reference data, the neuronal tissues within the white matter had the same conductance values. However, it was divided to show that separate structures could be expressed instead of multilayer modeling, and that the physical properties of each tissue could be adjusted. In future studies, the electrical properties, which are affected by the frequency of the tTIS, must be modified based on sophisticated measurements. For the electrode system, 69 electrodes were used, based on the international 10–10 system, which divides the reference points on the scalp into 10% increments, and the diameter of each electrode was set to 1 cm (Figure 3(c)).

**Table 1.** Electrical conductivity of organs.

| Material | Conductivity (S/m) | Material | Conductivity (S/m) | Material | Conductivity (S/m) | Material | Conductivity (S/m) |
| --- | --- | --- | --- | --- | --- | --- | --- |
| Scalp | 0.43 | Corpus callosum | 0.12 | Amygdala | 0.32 | Caudate nucleus | 0.32 |
| Skull | 0.015 | Hind brain | 0.25 | Hippocampal formation | 0.32 | Unclassified | 0.32 |
| CSF | 1.79 | Putamen | 0.32 | Nucleus accumbens | 0.32 |  |  |
| Grey matter | 0.32 | Thalamus | 0.32 | Globus pallidus | 0.32 |  |  |
| White matter | 0.15 | Third ventricle | 0.32 | Hypothalamus | 0.32 |  |  |

### 2.3 Stimulus formulation

Formulating the modulation generated by an electrode montage is necessary for optimizing and analyzing the results [2, 23]. The electric field for the electrode montage can be obtained as a linear combination of (1) for the unit-current electric field of each electrode with one reference electrode as the cathode.

In this study, it was assumed that modulation was entirely determined by the magnitude of the beats. The sensitivity of the target tissue to an electric field differs and is known to exhibit a nonlinear relationship [9]. The function of the relationship was modeled for only a few tissues, and many factors require verification, such as the connection between tissues through nerve fibers. These issues should be addressed in further studies and were not addressed in this study.

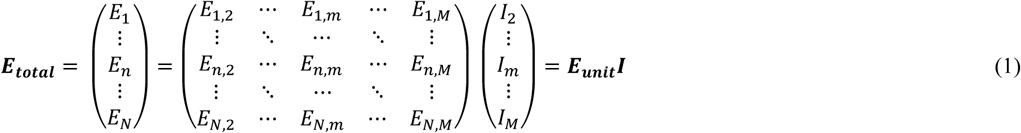

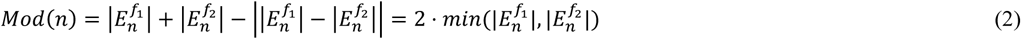

In this study, the magnitude of the interference modulation, which is assumed to be fully represented by the amplitude of the beats, is determined according to (2) as the absolute difference between the electric field amplitudes at two distinct frequencies. The beat frequency, on the other hand, corresponds to the absolute difference between the two frequencies, a relation that has been established in previous studies and adopted as the target frequency in tTIS. [2, 23, 28, 43]. In these equations, *m* and *n* represent the indices of the electrode and position in the head, respectively, while *M* and *N* denote the total number of electrodes and head positions. The amplitude refers to the zero-to-peak size of the sinusoidal signal, and f_1 and f_2 indicate two distinct frequencies. The distribution of the electric field is the negative gradient of the electric potential V, and the electric potential of each unit current for each electrode is calculated as a linear approximation using FEM, based on the Laplace equation [27]. The FEM of a realistic head model composed of tetrahedrons is implemented in a widely used method for brain stimulation [2, 23, 27, 28], and is written in FORTRAN.

### 2.4 Electrode montage optimization

#### 2.4.1 Evaluating Function

The distribution of modulation by the electrode montage was evaluated using the following three indicators: peak ratio (PR), concentration ratio (CR), and restriction ratio (RR) [2]. PR indicates the comparison of the peak modulation of the target and non-target, and CR evaluates the concentration of the spatial integral of the modulation on the target. Finally, RR represents a ratio controlled to have a value less than the average modulation of the target (AT) among the non-target parts.

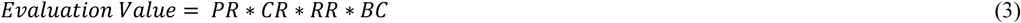

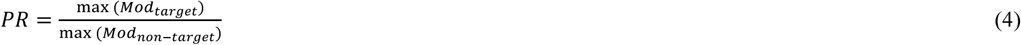

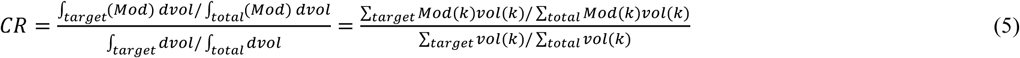

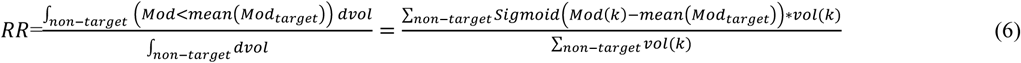

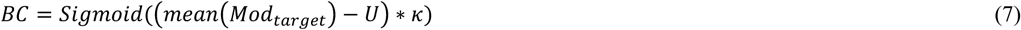

Optimization algorithms follow an iterative process of finding optimum solutions and require a single overall evaluation value to rate the solutions. Three ratios, PR, CR, and RR, which should be optimally as high as possible, were multiplied to provide an overall assessment. In addition to the product of indicators, the blocking condition (BC) prevents the magnitude of AT from falling below a specific value of U = 0.25 V/m [41, 43] when the highest current through the electrodes is normalized to 2 mA [23, 25]. Because all the evaluation factors are relative ratios, the magnitude of the modulation can be small compared with the injected current without these conditions. This value, as a threshold stimulus, is controversial from a biological perspective [40, 57]. However, this issue is not addressed here and is used only as a lower limit to prevent the amplitude from falling below a certain value. Parameter κ is a constant that adjusts the sharpness of the sigmoid function. This value should be sufficiently large to ensure the continuity of the evaluation value within the precision of the software; however, a smaller value ensures fewer artifacts of evaluation values.

#### 2.4.2 Optimization Algorithm

We propose a method to find an electrode montage that can stimulate the target region optimally, including an efficient electrode arrangement and electrode current, using PGA. This is a method that improves computational performance by inducing the expression to evaluating all objects in parallel when solving tTIS with GA. In addition to comparing the speed with traditional GA, we also verified the effectiveness of the electrode position optimization by comparing the optimization results of the high-definition electrode system with USNN.

A USNN can be used to obtain the current value of a high-definition electrode system to stimulate a target [2]. This network structure uses a stimulus formula network fixed to a generating network. The generating network generates an electrode current through the hidden layer from a constant unit. The loss function returns the reciprocal of the evaluation value of (3) and optimizes the generative network to minimize it. In this network, ReLU [56] was used as the activation function. Layer normalization [1] was performed between each layer, and the optimizer used was Adam [44] with a learning rate of 0.01 and decay of 0.001. The network structure and learning parameters were empirically designed through repeated testing. In this method, the electrode current can be quickly optimized based on the gradient descent with the inertia method using multiple hidden layers. However, this method cannot address the problem of electrode positioning in the 10–10 system, where the solution space is discontinuous. The USNN method can only be applied when the electrode position is fixed.

A GA is an optimization algorithm that obtains solutions by imitating the evolutionary process of living things [21]. This is a stochastic method in which solutions can be obtained, even in a discontinuous solution space [49]. Recent studies have attempted to optimize the tTIS through GA [54]; however, the calculation time is considerably long. Herein, we introduce the PGA to describe the construction of chromosomes to solve the tTIS problem and discuss the derivation of the matrix operation for computing acceleration using a GPU.

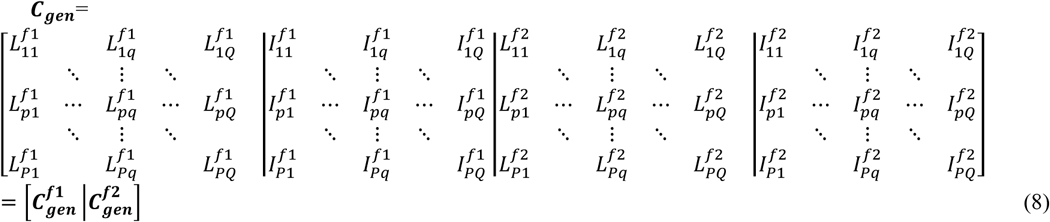

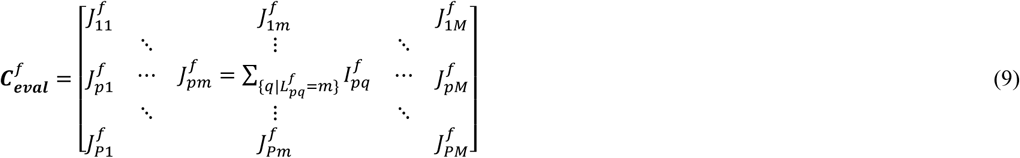

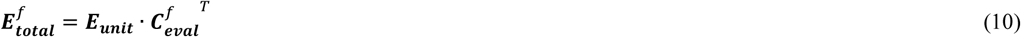

The chromosome composition is determined by the arrangement of the position index L of each electrode and current value I of the electrode in each row of (8). To ensure that the sum of the currents for each frequency in each individual row is zero, the value of the last Q-th current is determined as the negative value of the sum of all other current values without performing genetic calculations. This chromosomal arrangement, C_gen, performs genetic calculations of crossovers and mutations for all elements without encoding, where p is the individual index within the total population (P), and q is the electrode index within the total number of electrodes used for stimulation (Q).

For some special problems solved by GAs, the evaluation of objects can be processed simultaneously using GPUs, which has been shown to speed up computation [39]. Parallel processing was performed using a matrix operation to calculate the evaluation values for all chromosomes. For the matrix operation, C_gen is converted to C_eval, which is the current matrix for all electrode position candidates. J is the magnitude of the current through the electrode at position index m among the M electrodes (the total number of electrode positions). The products of C_eval and E_unit in (10) form the electric field distribution matrix, E_total, for all P objects at N locations, with a matrix size of P×N. The process of obtaining the modulation distribution using (2) and the evaluation value vector using (3) from E_total for each frequency can be obtained rapidly and simultaneously using a GPU for the matrix operation.

The detailed genetic operations of the GA (e.g., crossover, mutation, and elitism) are implemented in a general manner. For detailed implementations, the roulette-wheel method was adopted as the selection method, and the two-point crossover method was adopted as the crossover method. For a population comprising 1,000 individuals, 50 of them were selected as elite, and the mutation ratio was set to 20%. Mutation refers to modifications of genes to random values at random positions of the mutation ratio among all genes in the population. Detailed parameters, including the number of elites and mutation rates, were tested iteratively to obtain good empirical values.

#### 2.4.3 Current Scaling

As the evaluation value in (4)–(6) is determined to be relative, the electrode current must be scaled. In this study, electrode currents were scaled with 0.25 V/m as a reference, which is the numerical standard for misstimulation used in previous studies [41–43]. Figure 4 shows the changes of ratio (non-target <0.25[V/m]) and ratio (target >0.25[V/m]) when the maximum value of the returned electrode current is scaled in the range of 0–2 mA. As the scale increases, more areas of the target are stimulated; however, non-target misstimulation also increases. Because both ratios must be considered, the intersection where both values were equal was selected, allowing for scaling that balances target stimuli and non-target avoidance. Because the evaluation value had a BC, the intersection point was always less than 2 mA.

**Fig. 4.**
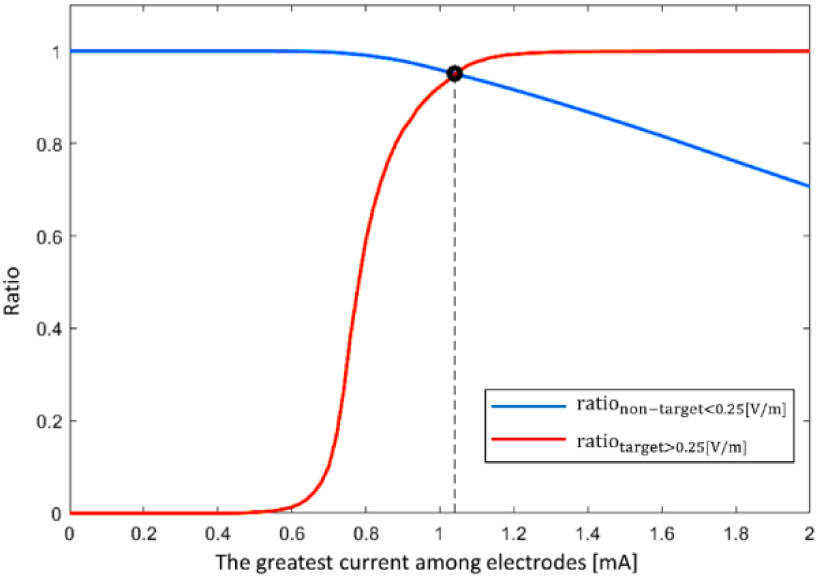
Intended volume ratio of the target and non-target according to the change in maximum current electrode.

### 2.5 Stimulation device

The stimulation device for tTIS should allow an alternating current with a carrier frequency of 2 kHz through the electrodes. Previous studies used devices that allow a given current to flow through any head using an isolator that converts the potential output to a current output. However, these methods do not function well at frequencies in the kHz range [10, 25, 31, 38]. Furthermore, despite the brain’s inability to respond to high-frequency noise, it necessarily requires a high precision voltage function generator to produce a clear voltage signal that the isolator can replicate through circuit feedback. In this work, we propose a method that utilizes the structure and properties of each tissue found in the previous step and the relationship between electrode current and internal head potential to directly find and apply a voltage output that produces the same result as applying an alternating current.

Because the external current flows only through the electrodes, the voltage generated at the electrodes by this current allows it to flow through them. The electrode voltage was calculated using the head model. In tTIS, the voltage applied to the alternating current flow to each electrode is not a simple sinusoidal wave but a combination of two sinusoidal wave frequencies. This complex waveform was generated digitally by PWM and applied to the electrodes through a low-pass filter and voltage follower (Figure 5(a)). PWM is a method of outputting a signal of desired magnitude by rapidly repeating an ON/OFF square wave signal with a duty cycle, which is the ratio of the ON signal to the OFF signal in a cycle [59]. This digital signal can be converted into an analog signal using a low-pass filter and voltage follower.

**Fig. 5.**
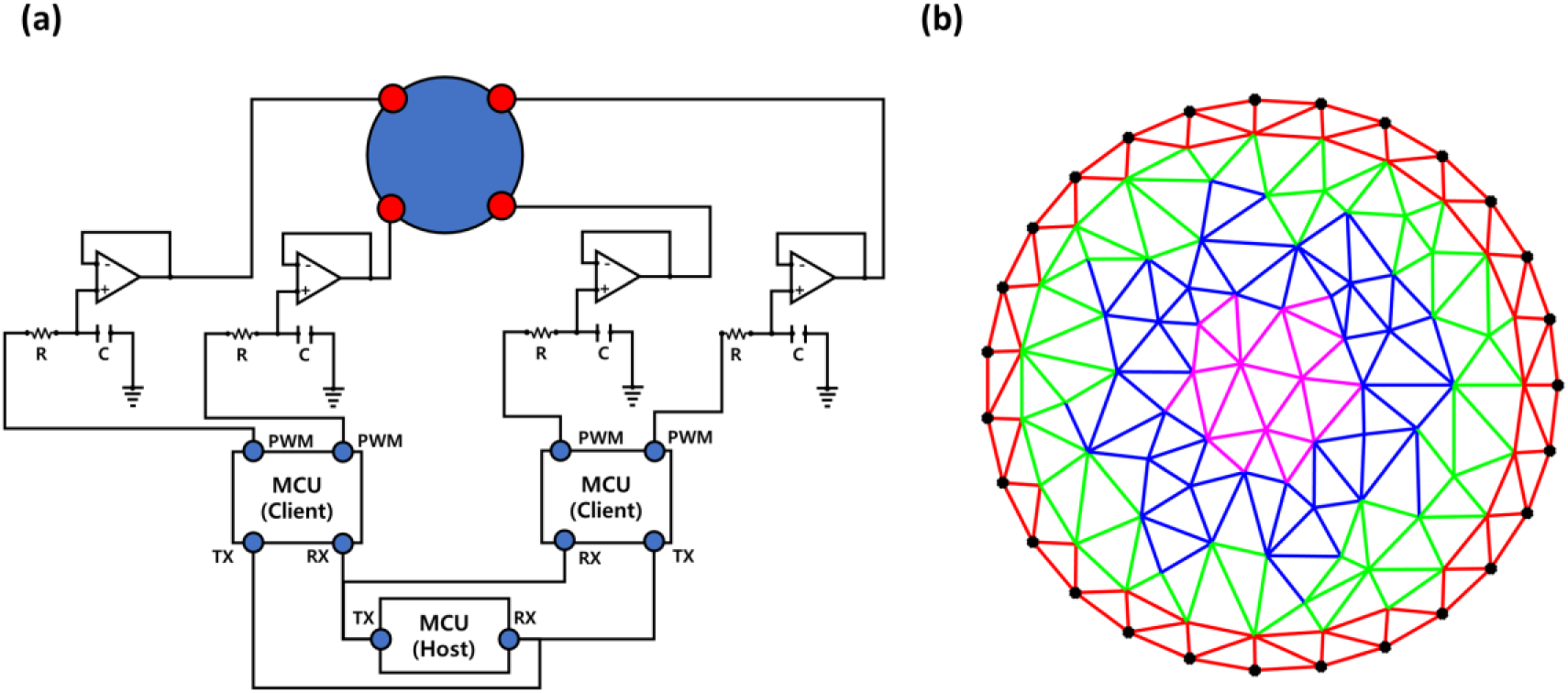
(a) Circuit diagram of the stimulation device (b) Schematic of the ideal head circuit.

The voltage output through the electrode is proportional to the duty cycle and is attenuated according to the frequency. This proportionality, which compensates for attenuation is measured in advance and applied once the frequency is determined.

The duty cycle function for the i-th electrode is as follows:

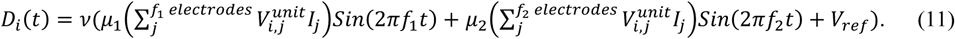

When unit current flows through the j-th electrode as an anode and reference electrode as a cathode, the voltage generated in the i-th electrode is V_(i,j)^unit, V_ref is the constant voltage of the reference electrode, ν is the duty cycle of the constant 1 V output voltage, and μ_1 and μ_2 are the amplitude attenuation ratios in a sine wave of frequencies f_1 and f_2, respectively. The current I_j through the j-th electrode is calculated in the optimization step. The duty cycle obtained in (11) was converted into a pulse using the Arduino library. The cut-off frequency of the low-pass filter, defined as 1/RC, was set to 10 kHz (R = 1 kΩ, C = 0.1 µF) to pass a sine wave of target frequency > 2,000 Hz and filter PWM frequencies of 100 kHz and high-frequency noise.

The MCU can generate duty cycles and PWM signals for many electrodes in each cycle. Therefore, as the number of electrodes increases, the MCU should be configured to handle only a specified number of electrode signals, as shown in Figure 5(a). In this experiment, an MCU was designed to calculate the PWM output of the four electrodes in a two-pair electrode set to determine the accuracy of the four-electrode setup for each MCU. Tx and Rx are the transmit and receive ports, respectively, for serial communication between boards.

### 2.6 Implementation details

#### 2.6.1 Computational Simulation Test

In this study, the optimization results were calculated and analyzed with the goal of stimulating the left thalamus of a realistic head model. Optimization was conducted for the electric field of the tetrahedral elements, such that spatial integration of the region of interest could be applied. The gray matter and deep-brain tissues were composed of 2,090,409 tetrahedrons (with 618,292 nodes), of which 100,000 tetrahedrons (with 278,137 nodes) were randomly selected for use. The original MRI model was imaged such that each voxel with an interval of 1 mm was downsampled to the expected spatial interval of 2,090,409/100,000 ^ 1/3, which was approximately 2.75 mm. This precision was used to calculate the evaluation value of the target and non-target, assuming that the electric field, calculated with 1 mm precision, was continuous throughout the head.

The test in the optimization process has the following objectives and processes. Initially, the evaluation focused on assessing the effectiveness of GAs in optimizing the electrode placement. This evaluation considered changes in the number of electrode pairs and generation to inform the selection of an appropriate electrode configuration plan for the clinical environment. In this test, two, three, four, and five electrodes per frequency were used based on the number of electrodes required for each frequency. This arrangement was expressed as two, three, four, and five pairs. We also analyzed the performance degradation of the optimal placement of fewer electrodes compared with the USNN results using all 10–10 system electrodes. Furthermore, we analyzed the computational time of the PGA method derived in this study and that of the conventional GA method.

In addition, to compare and analyze the effects of deep-brain modeling on the results, the optimization results were compared in the “uniform core model,” which has uniform conductivity in the white matter, and the “segmented core model,” in which detailed tissue was modeled. An electrode montage optimized for a model with a uniform core was applied to the head model with a segmented core. This verification process verifies how well an electrode montage, which works well in a uniform deep-brain model, stimulates a segmented deep-brain model, irrespective of the type of electrode system used. Therefore, this comparison was made using a three-pair GA that was selected arbitrarily.

#### 2.6.2 Ideal Head Circuit Test

The proposed tTIS stimulator must be validated because it uses an alternative method of finding and applying a potential input, which gives the same result as the current flowing through the electrodes. In addition, the use of digitally generated PWM signals in an alternative method can introduce noise and distortion. Therefore, it is important to verify the stability and reliability of the system. To verify the performance and accuracy of the hardware, excluding all artifacts generated when modeling the head, an ideal head circuit was used for the test. While the verification of the anatomical head in the stimulation situation was performed through 3D simulation, the ideal head model was only used to verify the stability of the hardware.

An ideal head circuit is constructed by mimicking the head using only resistors. It consists of a layer of high resistance (1,000 Ω: red), a layer of low resistance (220 Ω: green), and a set of layers of medium resistance (510 Ω: blue and magenta) in sequence from the outside (Figure 5(b)). To achieve stimulation, the blue resistor modulation should be minimized, and the magenta resistor modulation should be maximized.

The formula for the circuit analysis was organized in the same linear form as the FEM, using Ohm’s and Kirchhoff’s current laws [44]. The voltage difference between the nodes of any resistor is the product of the current and resistance, and the incoming and outgoing currents are the same at all the nodes. By combining these equations for all nodes, a linear system of equations for voltage and current can be obtained. The value corresponding to the electric field of the head is the potential difference mV/edge across the resistor. For the black dots in Figure 5(b), the nodes that form the outer boundary of the ideal head circuit are the candidate electrode nodes, and the electrode montage is optimized using a two-pair GA that selects two electrode positions for each frequency.

In this test, it was assumed that the amplitude of the beat determines the modulation. In this system, a sine wave with the same frequency and amplitude as the beats can be extracted through sampling. The sum of the two sine waves producing a beat at one location is

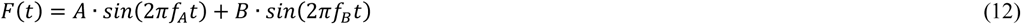

under the condition that |A|>|B|, the amplitude of the beat is 2×|B|, and the frequency is the target frequency of tTIS: |f_A-f_B|. If this signal is sampled with f_A—a time variable t=1/f_A (n+ɛ)—where n is a sampling number variable, and ɛ is a sampling offset greater than 0 and less than 1. Substituting this into (12) yields

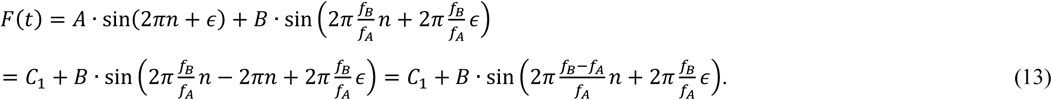

Substituting 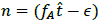 into (13):

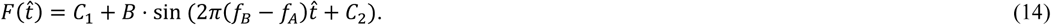

Here, C_1 and C_2 are the translation values unrelated to the frequency and amplitude, respectively.

If the signal of (12) is sampled with f_A, the amplitude component, A, becomes an offset constant, and a sine wave with amplitude B and frequency f_B-f_A is induced. This is a sine wave with the same frequency as the beats in (12), but with half the amplitude. The line tracing of the upper peak of the beats is slightly different from the sinusoidal form. However, the signal sampled by f_A becomes a sine wave by aliasing with the high-frequency beats. In this experiment, the PGA was used to determine the two-pair electrode positions and currents that could intensively stimulate the core of the ideal head model, which was then stimulated using the proposed stimulation method. During this stimulation, the potential difference across one edge of the target was measured, and the interference component was analyzed to determine how far it deviated from the expected result.

#### 2.6.3 Test Environment

All software development and computational testing were conducted on a system with an Intel-i9-9900k processor, 64 GB RAM, and an NVIDIA GeForce RTX 4090 GPU. The ground truth of the public network [4] was used for the segmentation of the realistic head model, and tetrahedral finite-element modeling was performed using the GIBBON Code library [33]. The FEM was written in FORTRAN in Visual Studio 2017, and the USNN and PGA were implemented using TensorFlow 2.3.0 and Keras 2.4.0, respectively, in Python 3.7.6. The microcontroller unit (MCU) for electrical stimulation used an Arduino Pro Portenta H7 board and was programmed using Arduino IDE 1.8.19. The connection of the circuit was implemented through a breadboard, and the model of the transistor used as an amplifier was an IC LM358P. Other analyses and calculations were performed using the MATLAB R2019b software.

## 3 RESULTS

### 3.1 Computational simulation test

Figure 6(a) shows a visual plot of the best solution among the 20 test results based on random seeds, with the highest evaluation value calculated using (3) for each setting and method. For the USNN, 2,000 epochs were executed, and for the PGA, 1,000 generations evolved. Box plots of the evaluation values of the results of the 20 tests for each optimization setting is shown in Figure 6(b). The resulting parameters, such as PR, CR, RR, and AT, for the best solution are listed in Table 2.

**Fig. 6.**
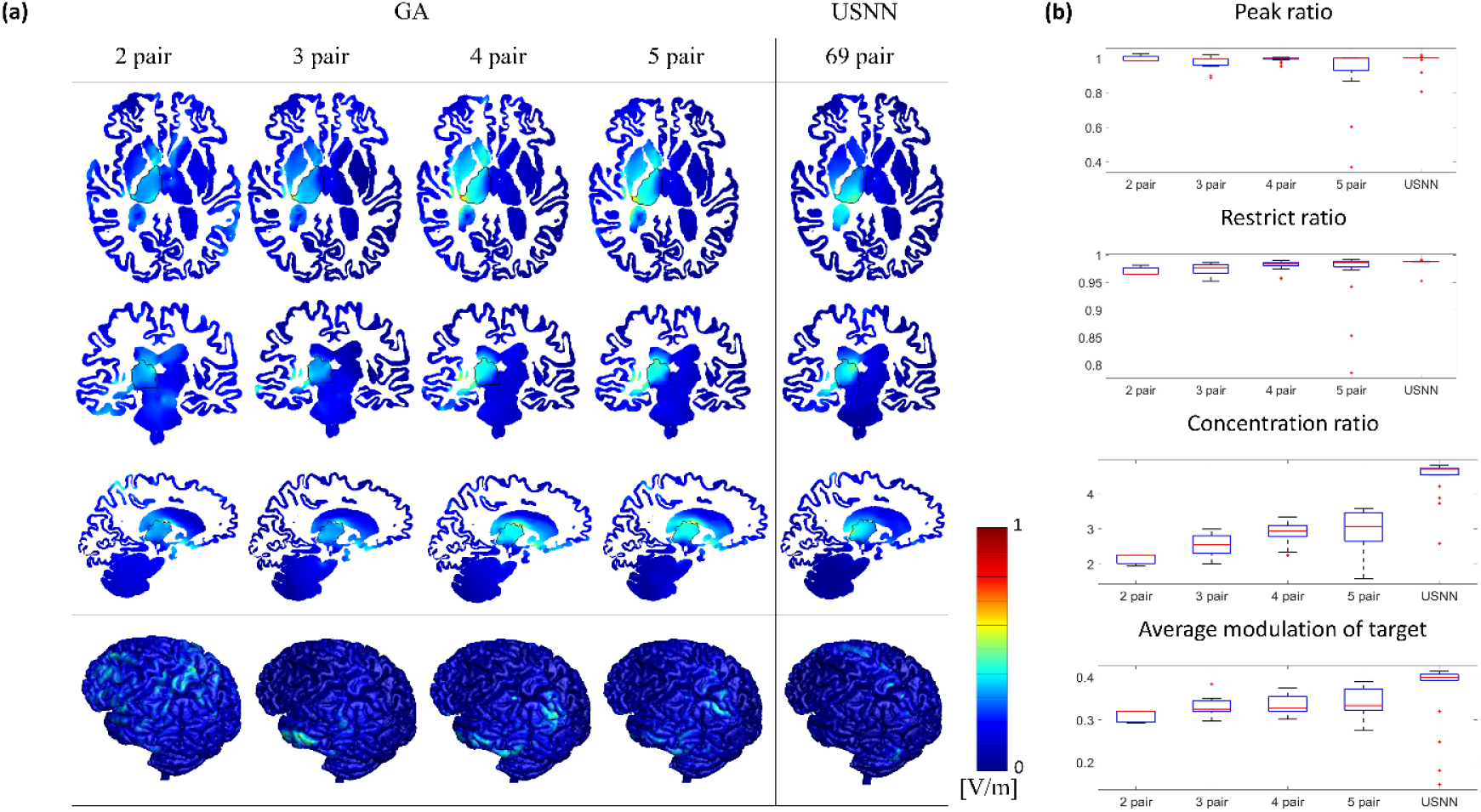
(a) Visual plot of the optimization results, (b) Box plot of the evaluation values for each optimization setting.

**Table 2:** Characteristics of the best solution among 20 results.

|  | GA |  |  |  | USNN |
| --- | --- | --- | --- | --- | --- |
|  | 2 Pair | 3 Pair | 4 Pair | 5 Pair | 10-10 System |
| PR | 0.9873 | 0.9994 | 1.0014 | 1.0017 | 1.0035 |
| CR | 2.2572 | 2.8412 | 3.3356 | 3.5909 | 4.7268 |
| RR | 0.9649 | 0.9871 | 0.9846 | 0.9866 | 0.9877 |
| AT [V/m] | 0.3199 | 0.3462 | 0.3738 | 0.3900 | 0.4115 |

In the case of the proposed PGA, 1,000 generations with a population consisting of 1,000 individuals were repeated in approximately 1 min, regardless of the number of pairs. In the case of USNN, it took approximately 8 min for 2,000 epochs. The conventional GA, which repeatedly evaluates all individuals, took approximately 2 h, despite the GPU calculation for each individual evaluation. Figure 7 shows the average evaluation value of 20 results per generation for each pair in the PGA.

**Fig. 7.**
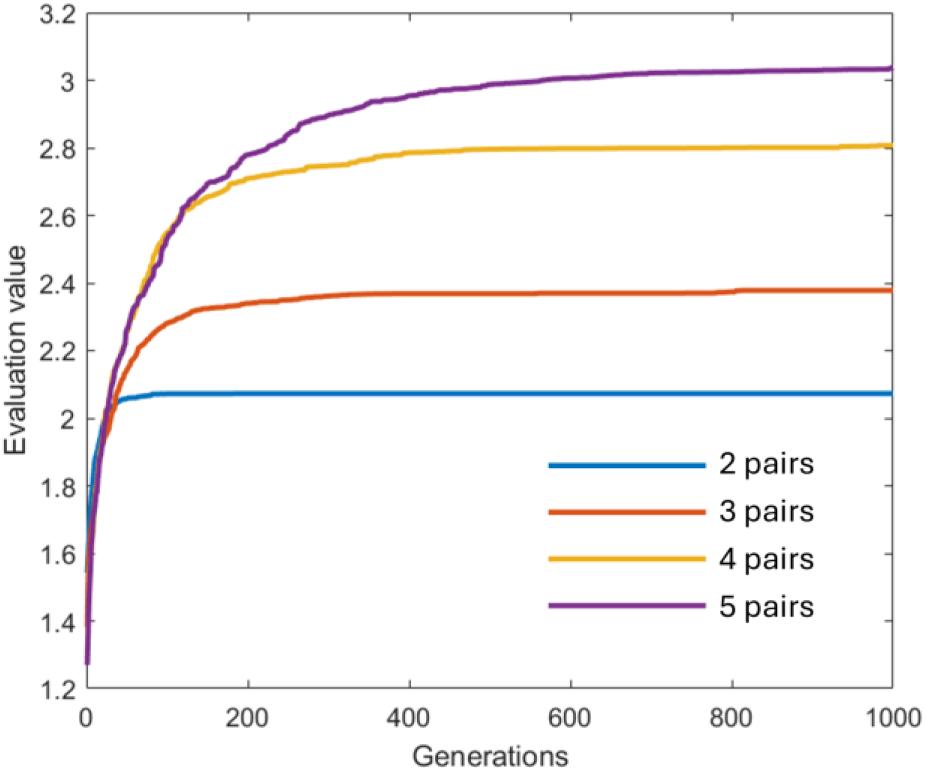
Average evaluation value of 20 results per generation for each pair in GA.

### 3.2 Comparative analysis of the realistic head model

Figure 8 shows the difference in visual plots according to the presence or absence of deep-brain modeling. Figure 8(a) shows the result of optimizing the uniform core head model with three pairs of GA, and Figure 8(b) shows the result of applying the same electrode montage to the segmented core head model. The electrode montage was optimized with evaluation values of 0.942, 3.074, 0.988, and 0.3072 V/m for the PR, CR, RR, and AT, respectively, for a uniform core head. However, the evaluation values decreased to 0.955, 2.514, 0.966, and 0.2394 V/m for the PR, CR, RR, and AT, respectively.

**Fig. 8.**
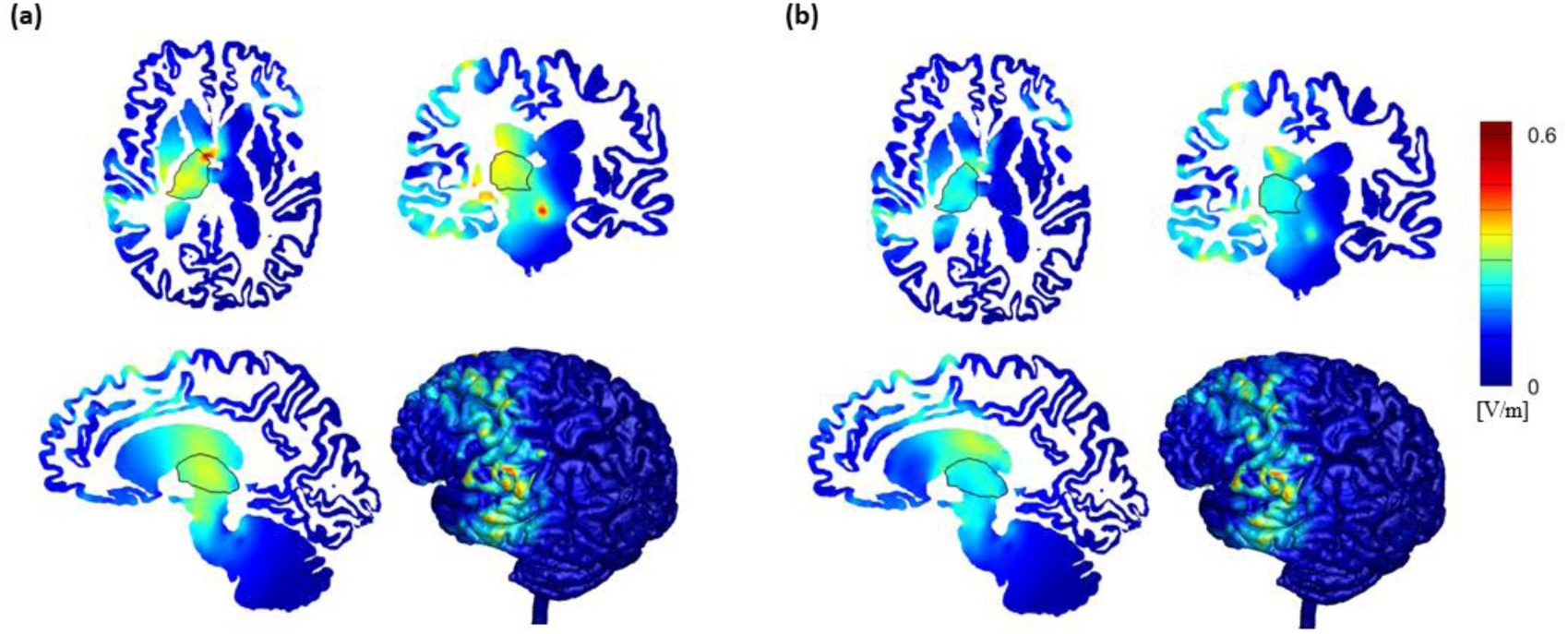
(a) Results of 3-pair optimization without modeling for deep tissues, and (b) Results of the same stimulation with deep tissues modeled.

### 3.3 Ideal head circuit test

Figure 9 shows a visual plot of the electrode montage and modulation optimized for the ideal head circuit. The evaluation value of the optimization was 3.8992 (PR: 1.445, CR: 1.575, RR: 0.857). When the maximum value of the electrode current amplitude is normalized to 1 mA, the amplitude of the current that must flow through the other electrode will be 0.5269 mA, a maximum modulation of 7.45 mV/Edge is obtained at one of the target positions.

**Fig. 9.**
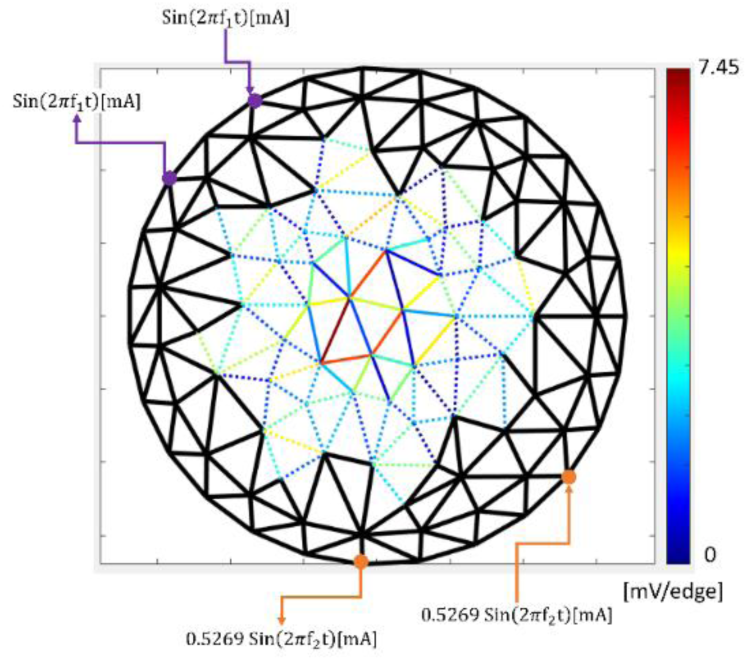
Electrode current and modulation results for the optimization of an ideal head circuit.

The duty cycle of each electrode was calculated using (11), based on the current obtained through optimization. Figure 10(a)–(e) shows the signal and frequency analysis for the edge with the greatest modulation in ideal head circuit optimization. Figure 10(b) shows the potential difference signal for this edge, confirming weak beat modulation with scrambled noise. Figure 10(c) shows the amplitude spectrum of the Fast Fourier transform (FFT) frequency analysis. It has high values at 2,000 and 2,020 Hz, and detectable noise at approximately 2,800 Hz. As the frequency of the beat is extremely low compared to that of the signal, it is difficult to verify it immediately from the FFT spectrum. However, at the beat of the edge, when A>B, by sampling the signal at f_A, Mod⋅sin(2π(|f_A-f_B |)t) can be obtained. The thin plots in Figure 10(d) show the sine waves of the modulation components of the 50 signals aligned by moving them parallel through template matching. The white line is the average of the data at each time step, and the cyan-colored line represents the actual potential difference calculated using the PGA and translated into least-squares error with the white line. The R^2 value, which is the coefficient for determining the actual potential difference for the average of 50 data points, is 98.21%. Figure 10(e) shows a boxplot for each frequency in the amplitude spectrum obtained by the FFT of each signal sampled at f_A.

**Fig. 10.**
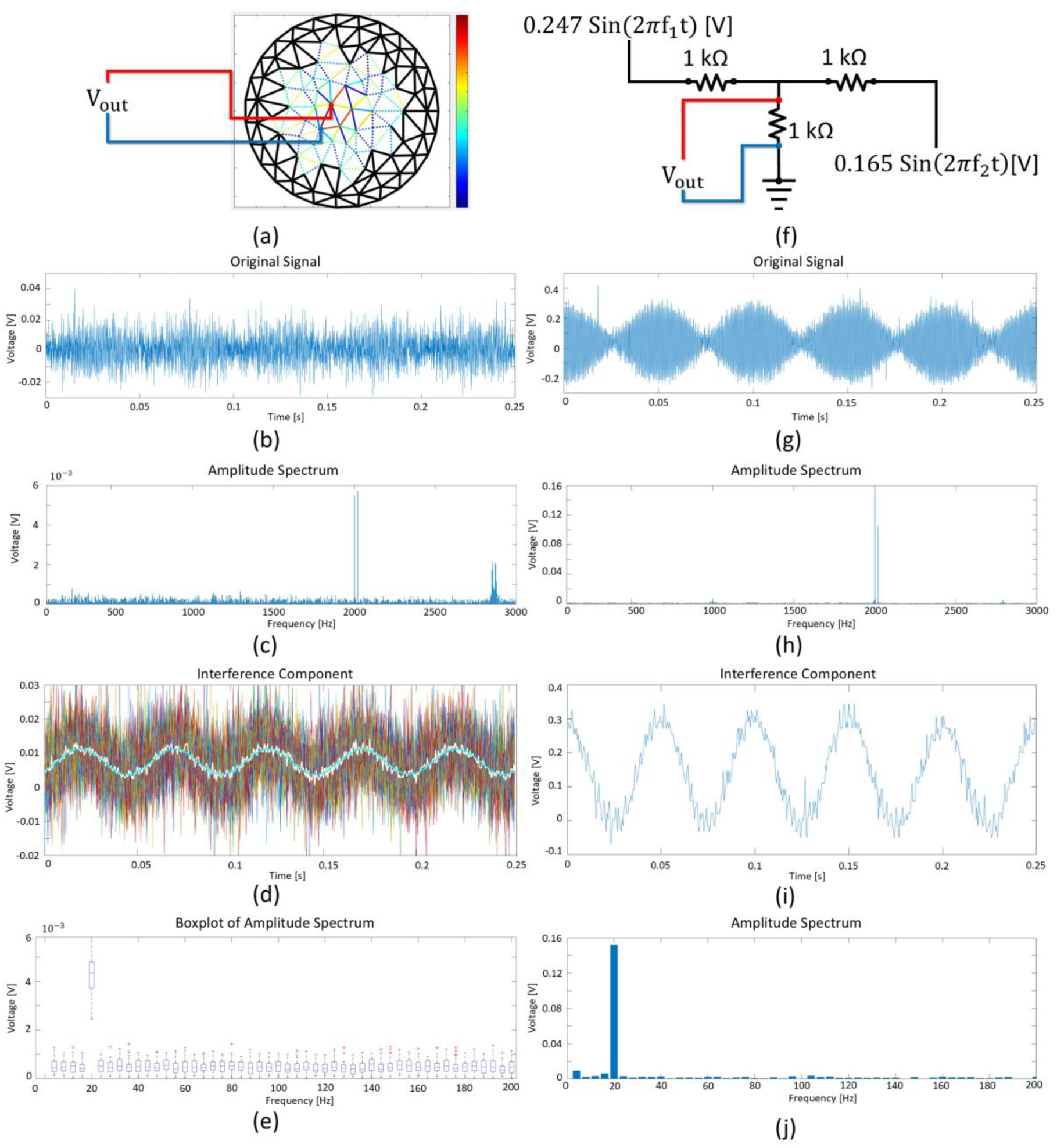
Results of potential difference measurement and frequency analysis of the target. (a) Measurement position in the ideal head circuit. (b) potential difference signal. (c) amplitude spectrum of (b). (d) interference component of (b). For thin-line results of 50 data points, the white line is the average value, and the cyan line is the predicted value. (e) Amplitude spectrum for each frequency in (d). (f) Simple test circuit for measuring interference signals excluding instrumentation noise. (g) –(j) Result corresponding to each (b) –(e) condition for the circuit (f).

The value of the current reaching the deep brain, despite the significant resistance layer, is exceedingly small, and the potential difference owing to the current is vulnerable to noise caused by the measurements. Additionally, to check the actual signal and noise of the board while minimizing the noise due to the measurement, a simple circuit that causes interference, as shown in Figure 10(f), was constructed. This test circuit was designed to check only the interference components of the signal via an unbranched and undistributed connection. Figure 10(g)–(j) represents the original signal, its FFT result, and the signal sampled with f_A and its FFT result of the simple circuit.

## 4 DISCUSSION AND CONCLUSION

In this study, an MRI-based precise multichannel tTIS system was designed for computing for an individual. The proposed system included processes that should be considered for realistic stimulation, from the stage of obtaining an MRI to the stimulation of the subject using a stimulation device. It was implemented by reflecting not only the issues of tTIS, that is, deep-brain modeling, nonlinear optimization for beats, and kilohertz stimulation, which did not need to be considered in the traditional tES, but also the issues pertaining to the entire clinical process. Moreover, segmentation through deep learning, finite-element modeling of segmented images, optimization through a GPU-based PGA, and digital signalization using a pre-interpreted head model were implemented in this system, and their effects and applicability were verified. These methods are suitable for solving tTIS problems, and can be applied to solve existing tTIS problems.

The computation was conducted on 3D voxel images, wherein detailed brain tissues were segmented through deep learning after the MRI image of the subject was captured. We introduced tetrahedral finite-element modeling to apply the FEM to the segmented 3D voxel image and evaluated the effects of deep brain modeling on the results. In previous tES studies that stimulated areas near the electrode, only the external parts of the brain were considered [27]. Moreover, various tTIS studies still used a brain model with a uniform core [2, 58]. However, stimulation performance using electrode montages, which ignored the complex structure of neuron-based organizations within white matter, fell short of expectations. In the three-pair electrode test, the decreased AT value was 0.2394 V/m, which was lower than the value of 0.25 V/m, which was the criterion of stimulation considered in the current scaling step.

Multichannel tTIS optimization requires large and complex calculations because the amplitude of the beats is a nonlinear combination of the amplitudes of two different frequency signals. A recent review of tTIS optimization research introduced two robust optimization computing methods. [60]. The first is to find the current of densely placed electrodes through USNN [2], and the other is to find the current and optimal placement of a small number of electrodes based on GA [58]. USNN can rapidly find well-optimized solutions, even in a solution space with a high dimensionality of 10–10 level electrodes. However, in clinical practice, attaching the 69 electrodes of a 10–10 system to the head and applying 138 sine waves are extremely difficult [26]. Although this is an efficient approach for accurately determining the positions of a small number of electrodes, it is a problem with a discontinuous solution space, and the USNN method, which assumes a continuous and differentiable solution space, cannot be applied to this discontinuous problem. In contrast, a GA can stochastically determine a solution through genetic operations, even in a discontinuous solution space [49].

As shown in the boxplot in Figure 6(b), the USNN with many electrodes yielded results with high evaluation values for all evaluation factors, and the results were stable even when using the high-definition electrode system. However, the PR, RR, and AT values of the best PGA results were not significantly inferior even when fewer electrodes were used. When using three or more pairs of electrodes, PR and RR showed less than 1% difference, and when using four or more pairs of electrodes, the AT reduction was lower than 0.0377 V/m (9.16%).

However, the CR was 24.03% lower than that of the USNN, even for five pairs. A low CR with a high PR and RR indicates that unwanted beats were generated throughout the brain in non-threatening magnitudes. This phenomenon can adversely affect brain stimulation, and it has been shown that the USNN has better focused stimulation performance. However, in terms of efficiency, in the case of a 10–10 electrode system using 138 sine waves at 69 electrodes, the increments in CR per electrode were extremely low. The PGA can effectively find an electrode montage with a convincing result through positional optimization with a few electrodes, resulting in significant savings in cost and effort when stimulating clinical situations, apart from the computation time. Increasing the number of electrodes is still a challenge in clinical settings, with issues including the application time and cost of equipment, subject fatigue, and accuracy of attachment [26].

The application of the GA to tTIS optimization requires heavy computation [54]. In general, the GA must perform iterations to evaluate a considerable number of populations in each generation, which requires numerous evaluation function calls. As shown in Figure 7, as the number of electrodes used increases, many generations are required for convergence, which is a major weakness of multichannel optimization. To evolve 1,000 generations with a population of 1,000 individuals, the traditional GA takes approximately 2 h, even if the GPU calculation same as USNN is performed for each individual evaluation. However, the PGA for the tTIS is suitable for parallel computation using a GPU to simultaneously evaluate all individuals while obtaining the same results as the convectional GA. In the evolution of 1,000 generations of a population of 1,000 individuals, PGA can be processed in approximately 1 min. Reducing the calculation time from the hour level to 1 min means that many subjects can have an MRI and then be stimulated on the same day, with no calculation day and no return visits. As shown in Figure 7, up to five pairs converged on average at 1,000 generations. However, as shown in the boxplot in Figure 6(b), the five pairs exhibited unstable results when the error bar was stretched down. Therefore, additional generations and several trials may be required, depending on the number of electrodes used, and the level of calculation can be adjusted by considering the waiting time from obtaining FEM results to stimulation.

Because tTIS uses a frequency in the kilohertz range as a carrier frequency that does not affect the brain cells [18], using a traditional device for tACS is difficult. Most commercial brain electrical stimulation devices [5, 35, 36] do not support kilohertz-level frequencies, and an isolator that converts the output of the function generator into current suffers from attenuation and distortion at that frequency [10]. Recently, an additional circuit was developed to solve this problem in the isolator, and it was verified that the voltage signal of the function generator can be reliably converted to a current signal [15]. Additionally, there are precise current-signal generators if the device is not considered heavy in multichannel situations. However, increasing the number of precise function generators to implement multichannels and connect isolators results in heavy and inefficient systems.

The proposed stimulation method utilizes the properties of all tissues and the relationship between the electrode current and the potential within the head to calculate and apply a voltage input that produces the same interference result by applying an alternating current at each electrode. Compared with the traditional isolator method, this method does not require additional equipment or troubleshooting techniques to convert the voltage signal from the function generator to a current signal by circuitous feedback. In addition, this relaxes the high level of noise control required by function generators for circuitous feedback and allows lighter circuitry and equipment. A special form of voltage signal that causes the same interference stimulus as the current source was implemented as a connection on the MCU chip. This has the advantage of being portable, programmable, and extensible for stimulation for clinical purposes.

The problem of finding an interference signal that stimulates the target area can be solved in a precisely modeled 3D head model and verified through simulation. However, because hardware that uses digital signals requires precision and noise verification, it was evaluated using an ideal head circuit and oscilloscope. In this test, two pairs of interfering stimulations were precisely applied to the ideal head model, with a coefficient of determination of 98.21%. Using a simple circuit (Figure 10(f)) to exclude the measurement noise and highlight the interference components, the FFT results show that the target frequency can be strongly stimulated; however, signal noise also exists. This is owing to the topology of the prototype, the performance of the amplifier, and the computational power of the MCU, which can be improved through module improvements. This result indicates the case wherein one MCU outputs four electrodes (eight sine functions), and if the number of electrodes per MCU increases, noise and distortion become severe. When the number of electrodes increases, stability can be improved by using multiple MCU boards to control the distributed electrodes.

In this study, the tTIS problem was solved and verified by considering the entire process in a clinical setting. However, various problems must be considered and solved in practice to develop a precise tTIS system. In previous studies, the significance of simulations at low frequencies was verified in in vivo experiments [30, 46]; however, the time-dependent electromagnetic effects of the head require consideration when using high-frequency signals [9]. Accordingly, an FEM based on a time-differential equation is required to determine the stimulus equation. The material selection for the electrodes also affects the accuracy of the injected current at high frequencies [10, 53]. Furthermore, the proposed system was implemented to generate an MRI-based signal; however, this was verified only through simulations and circuit tests. The biophysics and connectivity of the neural circuits, which are factors affecting the cellular response to stimulation, were not considered in this study. The simulation assumed uniform and isotropic conductivity under quasi-static conditions and ignored detailed biological models, such as the current source inside the brain, soma, and axon. Therefore, additional verification is required for the application of this system in clinical practice. To this end, animal experiments should be performed in a future study, such as c-Fos staining verification, which was used to verify the previous tTIS test [18, 51], or SEEG, which can measure signals at step positions in depth by inserting a needle into the brain [7].

## ACKNOWLEDGMENTS

This research was supported by the KBRI Basic Research Program through the Korea Brain Research Institute funded by the Ministry of Science and ICT (24-BR-03-01 and 24-BR-05-01).

